# Conformational Shannon entropy of mRNA structures from force spectroscopy measurements predicts the efficiency of −1 programmed ribosomal frameshift stimulation

**DOI:** 10.1101/2020.12.12.422522

**Authors:** Matthew T.J. Halma, Dustin B. Ritchie, Michael T. Woodside

## Abstract

−1 Programmed ribosomal frameshifting (−1 PRF) is stimulated by structures in mRNA, but the factors determining −1 PRF efficiency are unclear. We show that −1 PRF efficiency varies directly with the conformational heterogeneity of the stimulatory structure, quantified as the Shannon entropy of the state occupancy, for a panel of stimulatory structures with efficiency ranging from 2–80%. The correlation is force-dependent and vanishes at forces above those applied by the ribosome. This work supports the hypothesis that heterogeneous conformational dynamics are a key factor in stimulating −1 PRF.

---

Proteins are synthesized in the cell by ribosomes, which read an RNA message in single-codon steps until reaching a stop codon. Since each codon consists of 3 nucleotides (nt), there are 3 reading frames for every mRNA. Normally mRNA is read in a single reading frame, but in programmed ribosomal frameshifting (PRF), the reading frame is deliberately shifted at a specific location in the message to generate an alternate gene product [1,2]. Such programmed shifts into the −1 frame are commonly found in viruses, which use −1 PRF to produce two different polypeptide chains in a defined ratio [3–6]. The expression ratio of the two polypeptides is tightly regulated because it has critical effects on essential aspects of viral function, and disrupting this ratio can inhibit propagation or invasiveness [7–12].

Frameshifting is triggered by specific features in the mRNA (Fig. 1A): a slippery sequence of form XXXYYYZ, at which −1 PRF occurs, and a stimulatory structure typically located 6–8 nt downstream, which is usually a pseudoknot (a tertiary structure consisting of intercalated stem-loops) but sometimes a hairpin [2,13–19]. It remains unclear which properties of the stimulatory structure determine −1 PRF efficiency, however, in part because the mechanism of −1 PRF is still incompletely understood despite significant strides in elucidating the key steps involved [20–23]. Previous work has had some success in linking −1 PRF efficiency to static properties of specific stimulatory structures, such as particular structural features or their thermodynamic and/or mechanical stability [24–31], as well as to ribosome structural dynamics and kinetics [23,32]. However, none of these links has proven able to provide a general framework predictive of −1 PRF efficiencies across a wide range of conditions.

**Figure 1:**
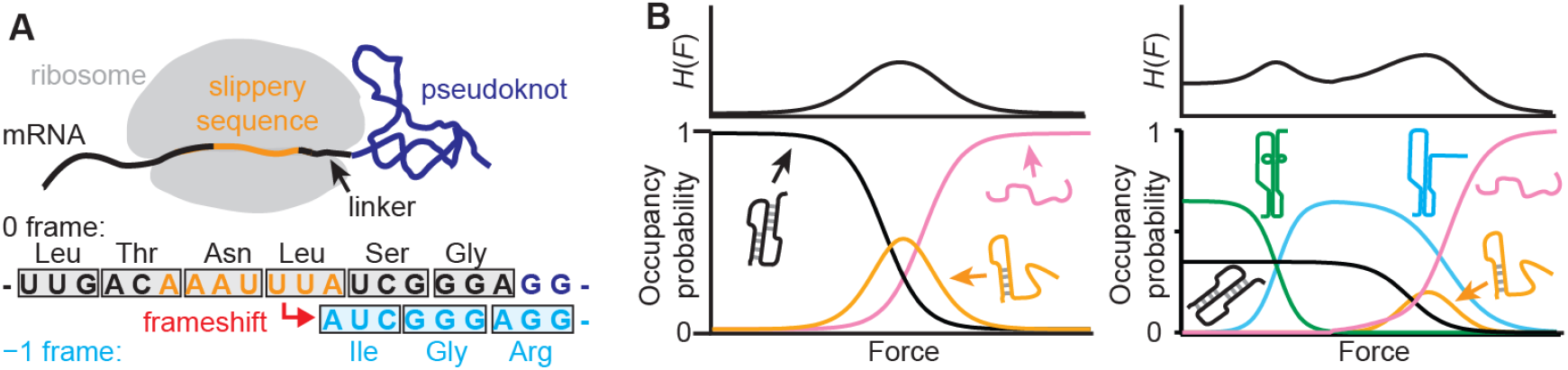
−1 PRF and conformational Shannon entropy. (A) −1 PRF is stimulated by a structure in the mRNA (here, a pseudoknot) just downstream of a slippery sequence, leading to a shift in the ribosome reading frame. (B) The Shannon entropy of the stimulatory structure as a function of force on the mRNA is illustrated notionally for a simple case where a pseudoknot unfolds via an intermediate (left) and a more complex case where multiple structures can be formed (right).

Previous work using optical tweezers [24,33–35] to study frameshift stimulatory structures while under mechanical tension—mimicking the situation during translation when the ribosome actively applies tension to the mRNA while resolving downstream structures [36]—found that high −1 PRF efficiency is linked with high conformational ‘plasticity’ or heterogeneity, reflected in the ability to populate more than one structure [33,37]. This link was found to hold true for a range of stimulatory structures, including both pseudoknots and hairpins [32,33,37–39], as well as when titrating anti-frameshifting ligands [40]. However, these studies were hampered by the lack of a physically meaningful metric of plasticity that can be applied under all conditions, raising questions about the meaning of the correlation and hence limiting the insight gained from it. Here we revisit the issue of conformational heterogeneity in frameshift stimulatory structures, introducing a new metric of heterogeneity that shows a strong linear correlation with frameshift efficiency. Not only does this new metric reconcile apparent contradictions between studies of wild-type versus mutant pseudoknots [24, 33,34], but it also shows a remarkable dependence on the tension in the RNA, with the strong linear correlation breaking down outside of the range of forces that are physiologically relevant in −1 PRF.

Motivated by the need to account for the possibility of multiple different states being occupied in a given force range as well as differences in the occupancies of these states, which may in turn be force-dependent, we quantified the conformational heterogeneity using the Shannon entropy [41,42]. Specifically, if at force *F* there are *N*(*F*) states occupied and state *i* has probability *P*_*i*_(*F*) of being occupied, then the conformational Shannon entropy is given by:

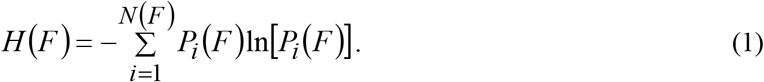

This measure is equivalent to the logarithm of the weighted geometric mean of the fractional state occupancies. When only a single state is occupied, *H* is minimal at 0, whereas *H* reaches a maximum of ln(*N*) when the *N* states are all equally likely to be occupied. *H* thus tends to increase as the number of conformational states available to the stimulatory structure increases (Fig. 1B).

We applied this new metric of conformational plasticity to measurements of the force-induced unfolding of different frameshift stimulatory structures. In each case, RNA containing the stimulatory structure being studied was attached via kilobase-long duplex handles to beads held in optical traps (Fig. 2A, inset). The traps were moved apart to ramp up the force and unfold any structure in the RNA, before relaxing the force to allow structures to refold and then repeating the unfolding force ramp. During each force ramp, we measured the end-to-end extension of the RNA constructs to report on changes in the RNA structure [35]. In the resulting force-extension curves (FECs), illustrated in Fig. 2A for two different frameshift-stimulatory structures, unfolding events appeared as sawtooth-shaped ‘rips’ where the length and force both changed abruptly. Different conformational states were distinguished in a FEC primarily through differences in the amount of unfolded RNA present, reflected in the unfolded RNA contour length (*L*_c_^u^) found by fitting each branch of the FEC to a worm-like chain polymer elasticity model [43] (Fig. 2A, dashed lines): each segment of a FEC involving a distinct *L*_c_^u^ value represented a different conformational state (Fig. S1A). Since different structures might in some cases share the same *L*_c_^u^, confounding an analysis based on *L*_c_^u^ alone, we also tested if the distribution of forces for unfolding states of given *L*_c_ ^u^ had the form characteristic for unfolding a single state [44] (Fig. S1B), and we tested if different folding pathways contained states with the same *L*_c_^u^, which would then necessarily be different (Fig. S1C). The unfolding forces also distinguished between structures with and without tertiary contacts, being characteristically smaller and more narrowly distributed for the latter (Fig. S1D).

**Figure 2:**
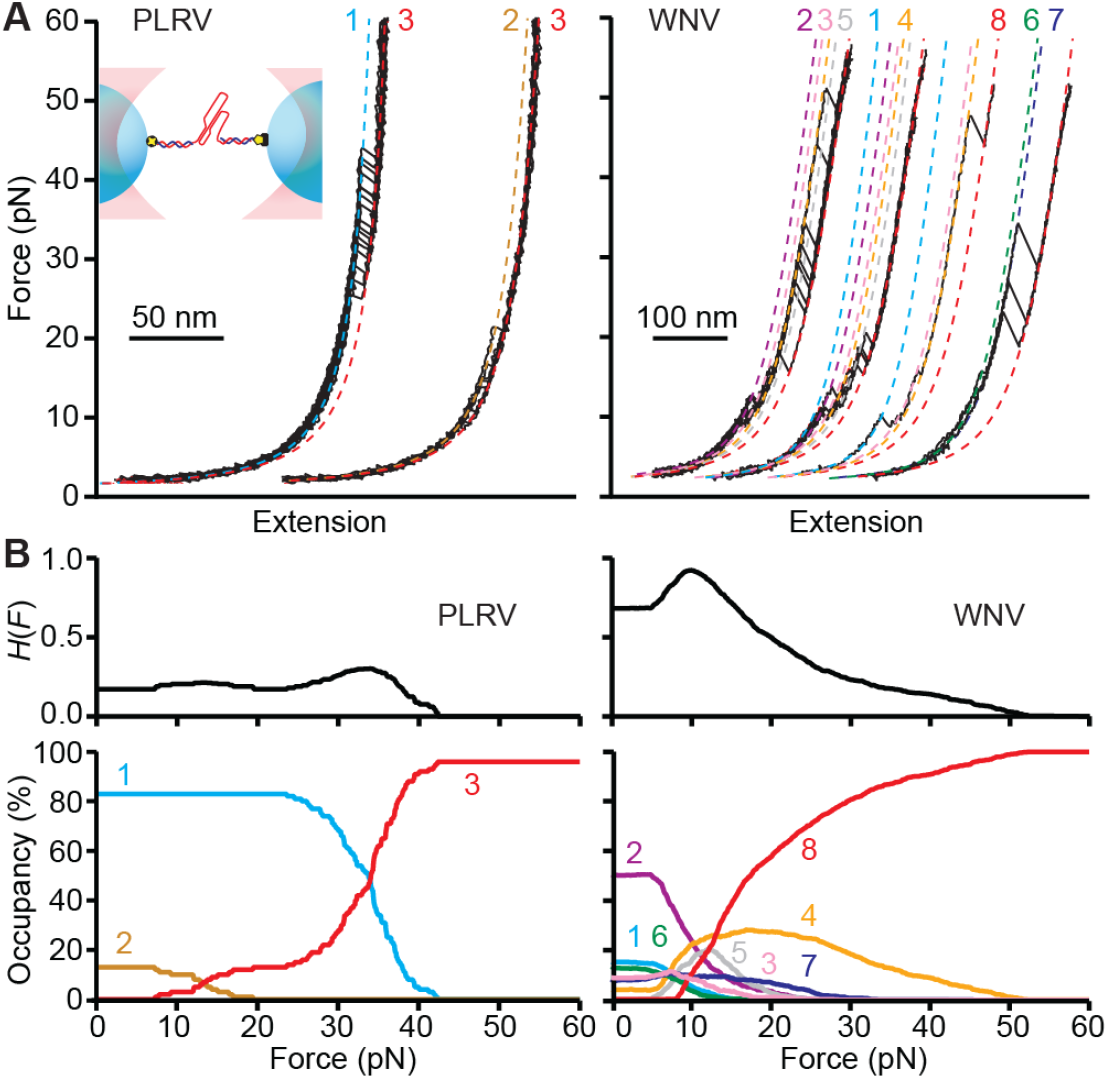
Force spectroscopy measurements of frameshift stimulatory structures. (A) Sample force extension curves of the stimulatory structures from PLRV (left) and WNV (right). For PLRV, 3 different states are seen (as numbered), whereas for WNV, 8 are seen. Sets of curves containing different states are offset for clarity. The states seen in stimulatory structures from different viruses are generally unrelated. Inset: schematic of measurement showing RNA connected by double-stranded handles to beads held in optical traps. (B) Fractional occupancy of each state as a function of force (lower panels) and force-dependent conformational Shannon entropy (upper panels) for PLRV (left) and WNV (right).

We converted each FEC for a given stimulatory structure into a force-dependent state trajectory, and then calculated the force-dependent probability of occupying each state based on the number of times it occurred in the collection of all FECs measured for that stimulatory structure (Fig. S2). The different states observed included the pseudoknots expected from each virus, the unfolded state, oftentimes one or more partially folded intermediates (*e*.*g*. hairpins) on the pathway to the pseudoknot, and in most cases one or more structures on alternative folding pathways (Fig. S1A). From the resulting force-dependent state occupancies for each of the stimulatory structures (Fig. 2B, lower panels), we then calculated *H*(*F*) via Eq. (1). The typical pattern for *H*(*F*) was that it started small to moderate at low force, where only the most stable states were occupied, it rose to a maximum at intermediate force as the force increased the relative stability of partially folded or alternative structures (thereby increasing the number of states occupied), and finally dropped to 0 at high force, where the RNA was fully unfolded (Fig. 2B, upper panels).

The calculation of *H*(*F*) was repeated for a large panel of stimulatory structures from different sources (Table S1), including the sugarcane yellow leaf virus (ScYLV), pea-enation mosaic virus-1 (PEMV), potato leafroll virus (PLRV), simian retrovirus-1 (SRV), human endogenous retrovirus-K10 (HERV), Visna-Maedi retrovirus (VMV), severe acute respiratory syndrome coronavirus-1 (SARS), West Nile virus (WNV), Middelburg virus (MIDV), mouse mammary tumor virus (MMTV), and human telomerase, as well as a non-stimulatory pseudoknot from phage T2 gene 32 (PT2G32) as a negative control. The measurements of all but the stimulatory structures from PLRV and MIDV have been reported previously [24,33,37]. Plotting *H* as a function of the −1 PRF efficiency, as reported for each structure from dual-luciferase assays [45], we found that there was a strong linear correlation between *H* and frameshifting efficiency at low forces (Fig. 3A, B), but that at forces higher than ∼20 pN this correlation vanished (Fig. 3C). Indeed, examining the force-dependence of the Pearson correlation coefficient, *r*, we found that it remained high for low forces until it dropped precipitously above ∼20 pN (Fig. 3D), crossing the critical Pearson coefficient for *p* = 0.01 (*r*_c_^2^ = 0.47, Fig. 3D cyan) at ∼23 pN. We verified that this correlation was robust against the outlying data at high −1 PRF efficiency by calculating the Spearman rank correlation (which is less sensitive to the effects of strong outliers), by recalculating the Pearson correlation without the data at high −1 PRF efficiency, and by comparing straight-line fits with and without the data at high −1 PRF efficiency (Fig. S3).

**Figure 3:**
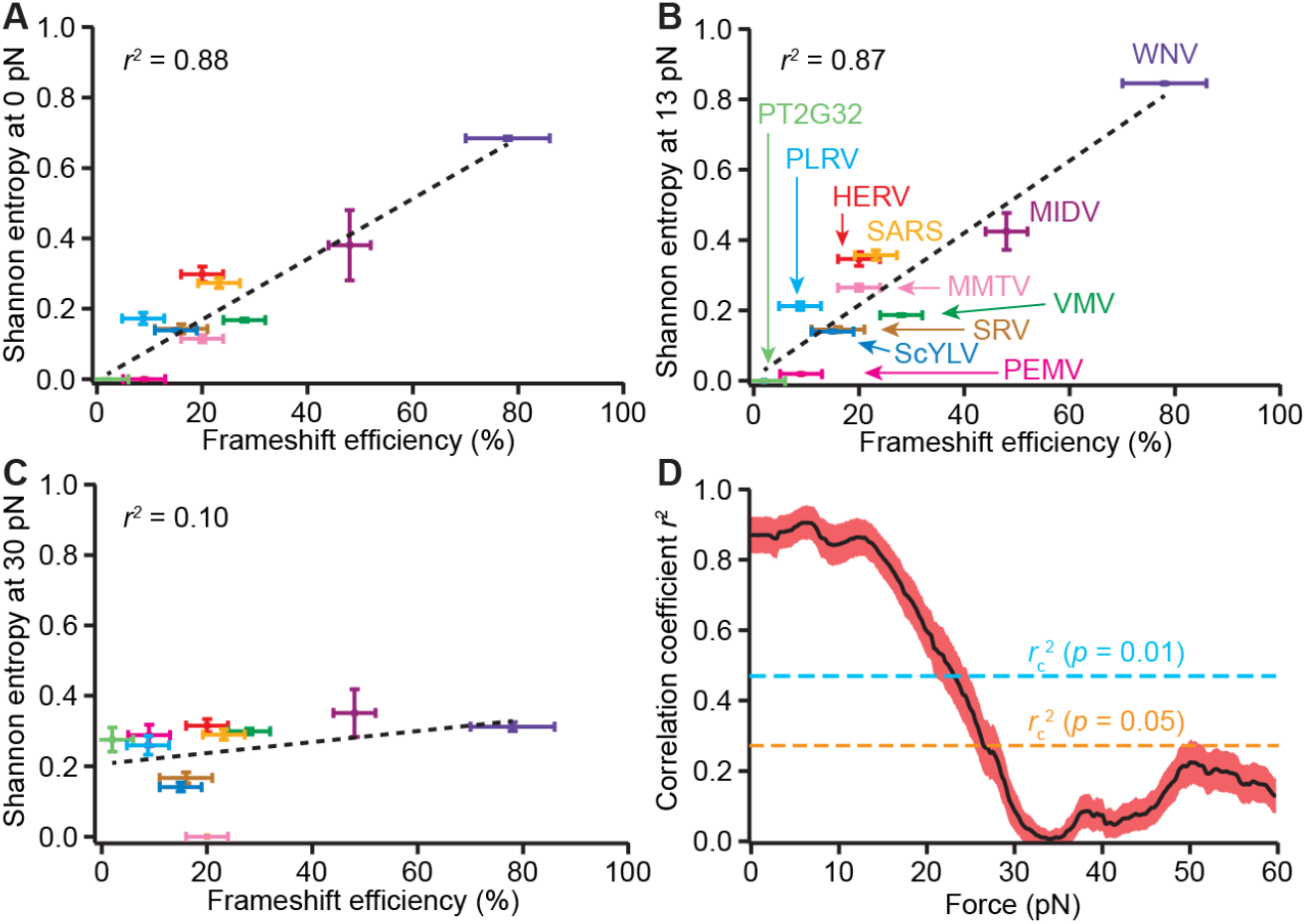
Correlation between Shannon entropy and −1 PRF efficiency for wild-type stimulatory structures. (A-C) Shannon entropy and −1 PRF efficiency vary linearly at low force (A: 0 pN, B: 13 pN) but are poorly correlated at high force (C: 30 pN) for the whole panel of wild-type stimulatory structures. (D) The Pearson correlation coefficient is strongly force-dependent, with the correlation vanishing above ∼20 pN, which is higher than the maximum force applied by the ribosome during translation (∼13 pN). Dashed lines show critical *r*^2^ value (cyan: 99% confidence level, orange: 95% confidence level), shaded region shows standard error in *r*^2^.

We next examined if the same correlation between *H*(*F*) and −1 PRF efficiency holds true when a stimulatory structure is mutated to suppress frameshifting. From measurements of four mutants of the human telomerase pseudoknot that stimulate different levels of frameshifting (Table S2), all reported previously [24], we calculated *H*(*F*) for each mutant and compared to the wild-type pseudoknot (Fig. 4A). We found a strong linear correlation in the force range ∼10–15 pN, although the correlation was absent for lower forces (Fig. 4B). Similar to the comparison between wild-type stimulatory structures described above, the correlation vanished at high forces, with *r*^2^ dropping below the critical value for *p* = 0.01 (*r*_c_^2^ = 0.87, Fig. 4B blue) above 15 pN. We note that these mutants were refolded without bringing the force all the way down to 0 pN, which may have biased *H* at low force, possibly accounting for the lack of correlation at 5–10 pN.

**Figure 4:**
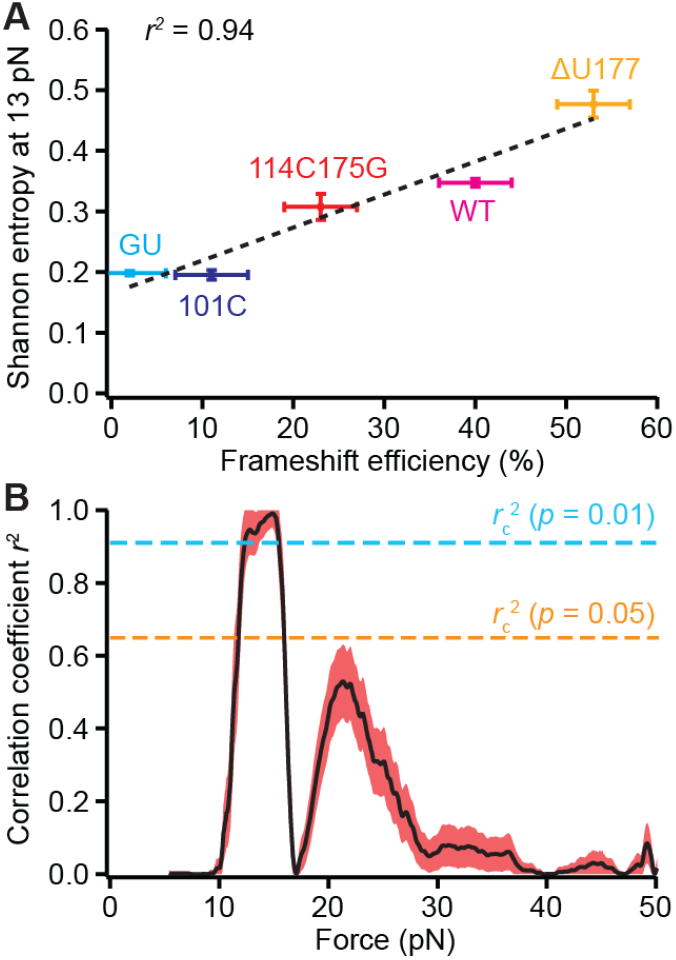
Correlation for mutant pseudoknots (color online). (A) Shannon entropy and −1 PRF efficiency vary linearly at 13 pN for mutants of the human telomerase pseudoknot. (B) The Pearson correlation again vanishes above the maximum biologically relevant force. Dashed lines show critical *r*^2^ values at 99% (cyan) and 95% (orange) confidence levels, shaded region shows standard error in *r*^2^.

Finally, we explored if the same correlation holds true when the −1 PRF efficiency is modulated by small-molecule ligands binding to the pseudoknot rather than changes in the RNA sequence. Small-molecule ligands that significantly alter the frameshifting efficiency have been explored as potential therapeutics for a number of viruses including HIV-1 [46,47], SARS-CoV [48,49], and SARS-CoV-2 [50,51]. We examined the effects of the binding of MTDB, a ligand that suppresses −1 PRF in SARS-CoV and SARS-CoV-2 [48,51] and was previously shown to reduce the conformational heterogeneity of the SARS-CoV pseudoknot [40]. Examining *H*(*F*) as a function of the ligand concentration for the data from Ref. [40], we were unable to compare it directly to −1 PRF efficiency, because MTDB appears to have weaker binding affinity to the pseudoknot in isolation in contrast to when it is complexed with the ribosomes during −1 PRF [40,48]. Using the fraction of pseudoknot that was bound by MTDB as a proxy for its effect on −1 PRF efficiency, however, we found again a linear correlation with *H*(*F*) for low forces, vanishing above ∼25 pN (Fig. 5).

**Figure 5:**
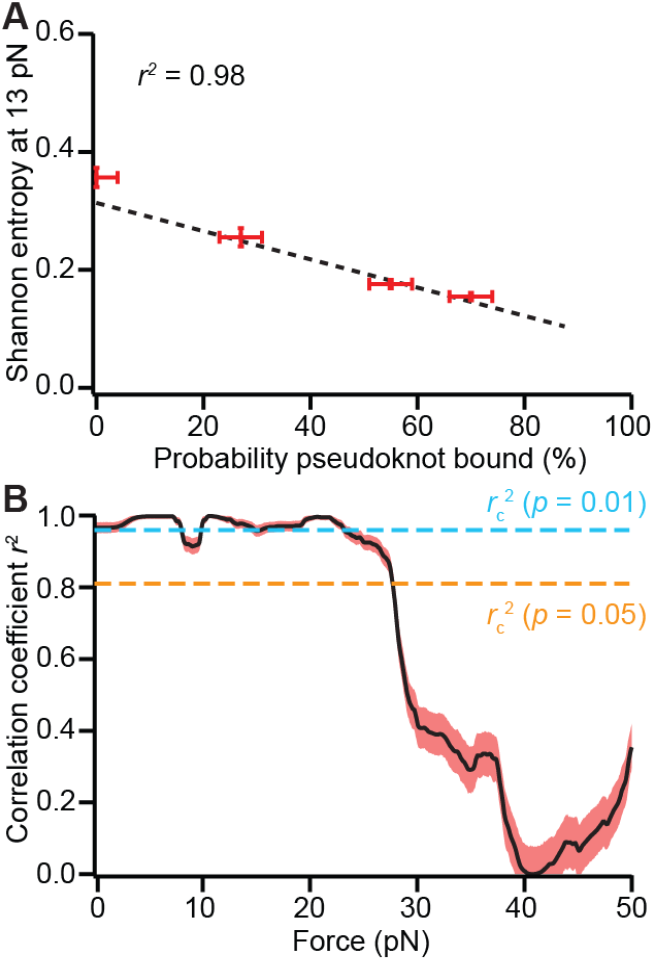
Correlation for pseudoknot bound with anti-frameshifting ligand. (A) Shannon entropy at 13 pN varies linearly with the fraction of SARS-CoV pseudoknot bound by a −1 PRF-inhibiting ligand. (B) The Pearson correlation vanishes above the maximum biologically relevant force. Dashed lines shows critical *r*^2^ values at 99% (cyan) and 95% (orange) confidence levels, shaded region shows standard error in *r*^2^.

The strong linear correlation between frameshift efficiency and *H*(*F*) observed both when comparing wild-type stimulatory structures and when comparing mutants with altered frameshift efficiency is an important result, because contradictory conclusions had previously been drawn from mechanical unfolding measurements of these two different sets of structures. Studies of the effects of mutating the nucleotides involved in base triples in pseudoknot structures found an exponential correlation between frameshift efficiency and unfolding force [24,34], but this result was contradicted by the lack of any correlation between frameshift efficiency and unfolding force when comparing different wild-type pseudoknots [33]. In contrast, the conformational Shannon entropy accounts for the frameshift efficiencies seen in both of these disparate datasets—in each case, the −1 PRF efficiency rises linearly with Shannon entropy—and even extends to the effects of frameshift-modulating ligands, suggesting that it provides a single explanation for all of the behavior observed, something that has not been possible until now.

The force-dependence of the correlation is particularly suggestive that *H*(*F*) captures an essential property driving −1 PRF, because the correlation is very strong in the range ∼10–15 pN but is gone for forces greater than ∼20–25 pN. Strikingly, this range matches closely to the range of forces applied by ribosomes translocating along mRNA: single ribosomes have been observed to exert up to 13 ± 2 pN before stalling [52]. Physically, the notion that the behavior of the stimulatory structure near the stall force of the ribosome is determinative of −1 PRF is reasonable: the ribosome is observed to pause at the stimulatory structure [23,32,53–55], ramping the tension up to the stall force multiple times as it attempts to unfold the RNA [22]. In contrast, one would not expect to see any correlation of the properties of the stimulatory structure with −1 PRF at high forces, beyond the ribosome stall force, because such forces are not physiologically relevant.

What insight does the success of the conformational Shannon entropy in predicting −1 PRF efficiency provide into the mechanism of frameshifting? The Shannon entropy depends not just on how many structural states are possible but especially on how often they are visited. Conceptually, it represents the uncertainty over what conformation is adopted, suggesting that it is the dynamical nature of the stimulatory structure—its ability to switch conformations—that is central to −1 PRF. This notion is in accord with previous work showing that static structures alone are insufficient to account for the extreme −1 PRF efficiency seen in some viruses [37]. The picture we propose is that changes in the conformation of the stimulatory structure induced by the ribosome as it forces the structure to unfold cause the tension in the mRNA to fluctuate abruptly; when these fluctuations are communicated to the mRNA-tRNA complex in the ribosome, the resulting disruption of the codon-anticodon interaction is guided by the slippery sequence to produce a shift into the −1 reading frame. This picture points directly at RNA stimulatory structure dynamics as a primary determinant of −1 PRF, complementing recently proposed mechanisms that link the induction of frameshifting to selective modulation of ribosome translocation dynamics by the stimulatory RNA [20–23,32,56]. Furthermore, we note this work has practical implications for efforts to target frameshift elements therapeutically by modulating −1 PRF efficiency: to inhibit −1 PRF, the aim should be to decrease *H* by increasing the stability of the most occupied state, whereas to enhance −1 PRF, it should be to increase *H* by stabilizing less-occupied states.

We note that this analysis of correlations between conformational Shannon entropy and −1 PRF efficiency has limitations. Discrepancies might arise from several sources. First, the optical trap measurements apply tension to both the 5′ and 3′ ends of the RNA, whereas the ribosome pulls on the 5′ end while pushing against the parts of the RNA structure abutting the mRNA entry pore. These differences in the geometry of force application might lead to somewhat different state occupancies; applying force with a nanopore [57] could improve the correlation. Second, although the state occupancies used to calculate *H*(*F*) are the non-equilibrium occupancies one would expect to be relevant based on the fact that the ribosome ramps the force out of equilibrium as it fluctuates over the slippery sequence [22], the ramp rates in the trapping measurements do not perfectly match those applied by the ribosome, which may alter *H*(*F*). Third, the trapping measurements were done in the absence of ribosomes, yet specific interactions between pseudoknots and ribosomes are thought to be important for modulating −1 PRF efficiency [29,31,58–62]. Hence it is likely that other factors that are not accounted for here also play a role in setting −1 PRF efficiency, which could be probed in future experiments using tweezers to observe the dynamic interactions of ribosomes with pseudoknots during −1 PRF. Nevertheless, as measured from the *r*^2^ value, the conformational Shannon entropy captures at least 85% of the variance in the −1 PRF efficiency in numerous stimulatory structures having a wide range of efficiencies modulated in multiple ways, suggesting that this metric incorporates the essential features determining −1 PRF efficiency. Future studies extending single-molecule observations of the dynamics of stimulatory structures as they interact with ribosomes during programmed frameshifting events [22,23,32] hold out promise for clarifying how the conformational dynamics of the stimulatory structures interact with ribosomal dynamics to induce frameshifts.

## Acknowledgements

We thank Prof. Gang Chen for generously sharing data for optical trapping measurements of the human telomerase pseudoknot. This work was funded by the Canadian Institutes for Health Research and the National Research Council of Canada.

