## Supplementary Information for "Conformational Shannon entropy of mRNA structures from force spectroscopy measurements predicts the efficiency of −1 programmed ribosomal frameshift stimulation"

**This PDF file includes**:

Supplementary Methods
Supplementary References
Tables S1 and S2
Figures S1–S4

**SUPPLEMENTARY METHODS**:

**Single-molecule force spectroscopy measurements**: RNA constructs for optical tweezers measurements were prepared as single transcripts containing roughly kilobase-long ‘handle’ sequences flanking on each end the sequence of the stimulatory structure of interest. Transcripts were annealed to single DNA strands complementary to the ‘handle’ regions, which were in turn attached to polystyrene beads held in optical traps. Optical trapping measurements were done as described previously [1,2]. Briefly, for the panel of stimulatory structures in Fig. 3, the traps were moved apart at 200 nm/s to ramp up the force, then brought back together to ramp the force back down to zero, holding the RNA near 0 pN for 3 s before repeating the cycle [1]. For the panel of mutants in Fig. 4, the traps were moved at 100 nm/s, cycling between ~3–5 pN and 60 pN [2].

**State identification**: The force-extension curves (FECs) measured for unfolding each stimulatory structure were analyzed to identify states as described previously [3]. Briefly, all segments of a given curve separated by ‘rips’ indicating structural transitions were fitted to extensible worm-like chain (WLC) models of the polymer elasticity [4] for the duplex handles and single-stranded unfolded RNA. Different structural states were distinguished primarily from differences in the contour length of unfolded RNA present in each case. In some cases, states with similar unfolded-RNA contour lengths were distinguished by their different unfolding forces or different unfolding pathways. WLC fitting parameters for the duplex handles were $L_p$ = 30–50 nm, $L_c$ ~ 900–1200 nm, and $K$ ~ 1200–2000 pN (depending on the construct). For the unfolded RNA, the WLC parameters were treated as fixed, at $L_p$ = 1 nm, $L_c$ = 0.59 nm/nt, and $K$ = 2000 pN, so that the only free parameter was the number of nucleotides of unfolded RNA in each segment of the FEC.

Note that the conformational Shannon entropy was calculated only for unfolding FECs. In principle, the same analysis could be applied to refolding FECs, but in practice it can be challenging to determine reliably the conformational state of the molecule during refolding FECs, because structures with similar $\Delta L_c$ but different bonding patterns (*e.g.*, simple stem-loops vs. more complex conformations with tertiary contacts) cannot be as readily distinguished from their refolding forces, whose differences are far less pronounced than for unfolding forces. We tested if the −1 PRF efficiency was correlated with the probability of refolding into any metastable structure from the (unfolded state) as a function of force, and found that there was no such correlation: the correlation coefficient was everywhere below the expected critical value at $p$ = 0.05 (Fig. S4).

**−1 PRF efficiencies**: Values for the −1 PRF efficiency for each stimulatory structure (listed in tables S1 and S2) were taken from the literature, using only those results measured in rabbit reticulocyte lysates to avoid possible systematic differences between the results from different assay types.

**Correlation significance tests**: To test the null hypothesis that there was no correlation, (*i.e.*, $r^2$ was not different from 0), we used the critical value for a one-tailed test at significance level $p$ = 0.01, except where otherwise indicated.

**Experimental uncertainties**: As experimental errors were not always reported in the literature for −1 PRF efficiencies, we conservatively estimated the error at a minimum of ± 4 percentage points, based on those studies that did report errors. The uncertainty in the Shannon entropy was calculated as

$\delta H = -\sum_{i=1}^{N(F)} \left[\ln P_i(F) + 1\right]\delta P_i$, where $\delta P_i = \sqrt{\frac{P_i(1-P_i)}{n}}$ is the standard error of a proportion.

**Table S1: Panel of stimulatory structures measured for Figure 3**.

| Name | NCBI Accession | Sequence | −1 PRF efficiency | Ref |
|---|---|---|---|---|
| PT2G32 | X12460.1 | UGACCAGCUAUGAGGUCAUACAUCGUCAUAGCAC | 2% | [5] |
| PEMV | MK948533.1 | AAUUCCGGUCGACUCCGGAGAAACAAAGUCAA | 9% | [6] |
| ScYLV | MF426270.1 | AAGUGGCGCCGACCACUUAAAAACACCGGA | 15% | [7] |
| SRV | 1E95_A | GCGGCCAGCUCCAGGCCGCCAAACAAUAUGGAGCAC | 16% | [8] |
| MMTV | M16766.1 | GGGGCAGUCCCCUAGCCCCACUCAAAAGGGGGAU | 20% | [9] |
| HERV | AB047209.1 | GGGGCCAGCCUCAGGCCCCACAACAAACUGGGGCAU | 20% | [5] |
| VMV | AAM51650.1 | AGGGGGCCACGUGUGGUGCCGUCCGCGCCCCCUAUGUUGUAACAGAAGCACCACC | 28% | [10] |
| SARS | AY291315 | GCGGUGUAAGUGCAGCCCGUCUUACACCGUGCGGCACAGGCACUAGUACUGAUGUCGUCUACAGGGCU | 23% | [11] |
| PLRV | MK116549.1 | UAGCGGCACCGUCCGCCAAAACAAACGG | 9% | [12] |
| MIDV | KM115530.1 | GUAAGCCUGGGAAUGGGGGCGACCCAGGCGUAUGAACAUAGUGUAACGCUCCC | 48% | [13] |
| WNV | NC_009942 | GGGCCUUCUGGUCGUGUUCUUGGCCACCCAGGAGGUCCUUCGCAAGAGGUGGACAGCCAAGAUCAGCAUGCCAGCUAUACUGAUUGCUCUGCUAGUCCUGGUGUUUGGGGG | 78% | [3] |

**Table S2: Panel of telomerase pseudoknot mutants studied in Figure 4**. Mutations shown in red text. From Ref 2.

| Mutant | Sequence | −1 PRF efficiency |
|---|---|---|
| WT | GGGCUGUUUUUCUCGCUGACUUUCAGCCCCAAACAAAAAAGUCAGCA | 40% |
| 114C/175G | GGGCUGUCUUUCUCGCUGACUUUCAGCCCCAAACAAAAGAGUCAGCA | 23% |
| ΔU177 | GGGCUGUUUUUCUCGCUGACUUUCAGCCCCAAACAAAAAAUGUCAGCA | 53% |
| GU | GGGCUGUUUUUCUCGCUGACUUUCAGCCCCAAACGUAAAAGUCAGCA | 2% |
| 101C | GGGCUGUUCUUCUCGCUGACUUUCAGCCCCAAACAAAAAAGUCAGCA | 11% |

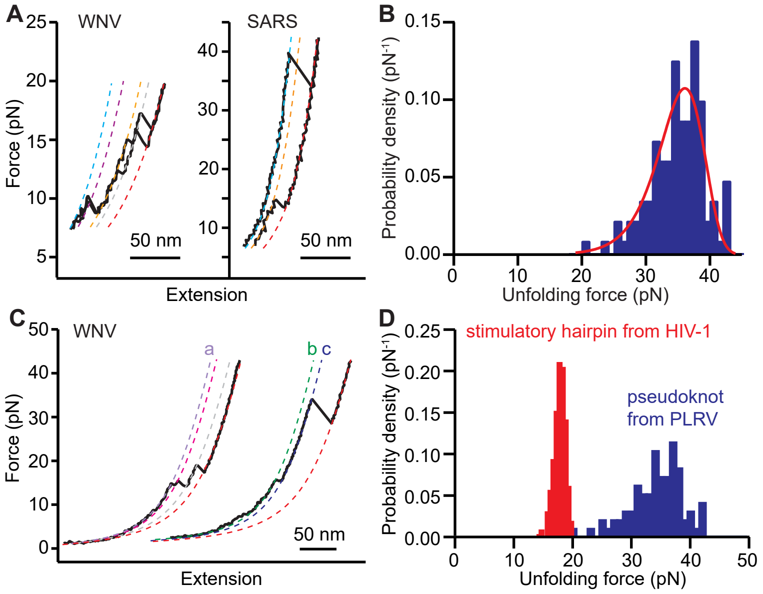


**Fig. S1: Identifying different conformational states**. (A) Segments of FECs that fit to WLCs with different $L_c^u$ (dashed lines) represent different conformational states. Right: the two FECs start in states with different $L_c^u$, indicating they are on different pathways. (B) The distribution of unfolding forces has a characteristic shape for unfolding a single conformational state; fitting to this shape (red line: fit to theory from Ref. 44) can test for the presence of multiple states with similar $L_c^u$ (here, the fit indicates unfolding of a single state). (C) If states with similar $L_c^u$ always appear in distinct pathways, they must represent different conformations. Here, state b has the same $L_c^u$ as state a, but only ever appears in combination with state c, whereas state a never appears in combination with state c, indicating they lie on difference pathways. Adapted from Ref. 37. (D) Unfolding forces for structures that contain no tertiary contacts are narrowly distributed and low (typically below 20 pN), as seen here for the stimulatory structure from HIV-1 (red), which is a simple hairpin. In contrast, unfolding forces for structures with tertiary contacts are higher and more broadly distributed, as illustrated for the pseudoknot from PRLV (blue).

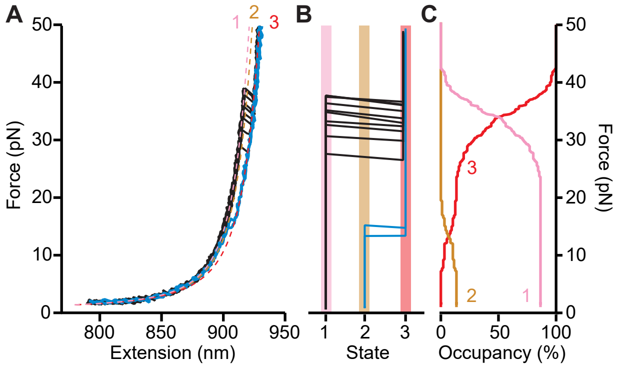


**Fig. S2: Analyzing state occupancies from force-extension curves**. (A) FECs were fitted to worm-like chain models (dashed lines) to identify states with different amounts of unfolded RNA and/or different unfolding forces. Here two different populations of FECs (black, blue) reveal three distinct states: a high-force pseudoknotted state (1, pink), a low-force state (2, brown), and the unfolded state (3, red). (B) Each FEC is transformed into a state trajectory, showing the state that is occupied as a function of the force on the RNA. (C) The fractional occupancy of each state as a function of force is found by averaging over all FECs.

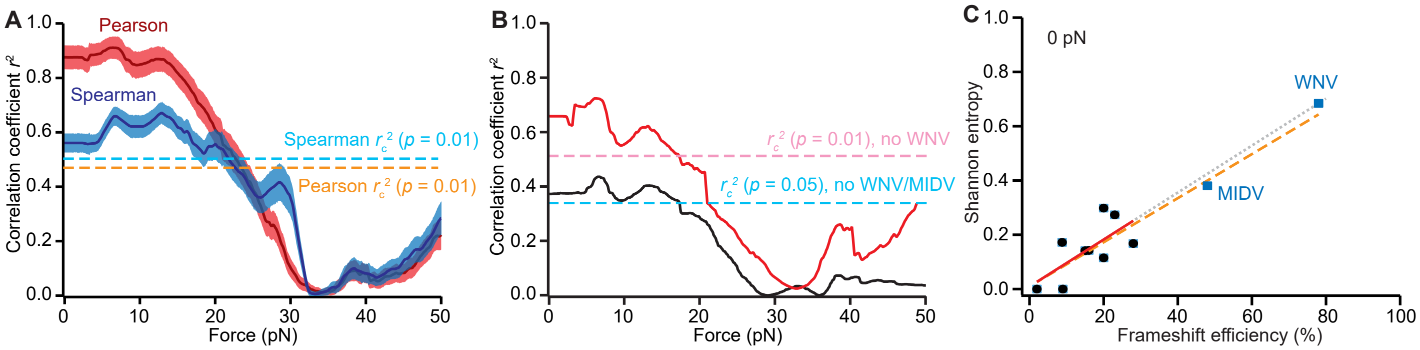


**Fig. S3: Robustness of correlation**. (A) The Pearson correlation from Fig 3D (red) was compared to the Spearman rank correlation for the same data (blue); both were significant at the 99% confidence level for forces up to just over 20 pN. (B) Testing the robustness of the Pearson correlation to removing the outlying data-points from WNV and MIDV, the correlation was significant at the 99% confidence level for force up to just under 20 pN when removing the data from WNV, and significant at the 95% confidence level when removing the data from both WNV and MIDV. (C) The straight-line fit (red) to the data for frameshift efficiency below 30% (black) at 0 pN is the same within error as the fit to all data (orange), and when extrapolated to high frameshift efficiencies (grey) predicts the observed values for MIDV and WNV (blue) very well.

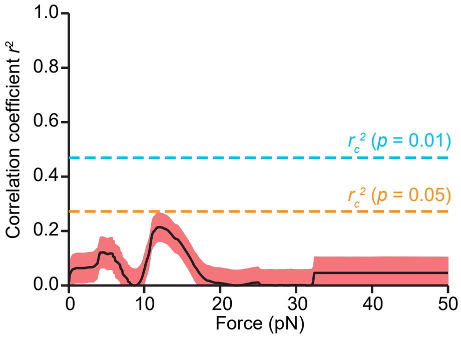


**Fig. S4: Frameshift efficiency is uncorrelated with refolding probability**. The Pearson correlation of the −1 PRF efficiency with the probability of refolding into any structure is below the critical value for the 95% confidence level at all forces.
